# In vivo imaging of reactive oxygen species after myocardial ischemia-reperfusion injury: a large animal multimodal imaging and transcriptomic study

**DOI:** 10.64898/2025.12.04.691257

**Authors:** Sophia Swago, Chiara Camillo, Marina Awad, Evan Gallagher, Elizabeth W. Thompson, Estibaliz Castillero, Ting Peng, Liming Pei, Zhiliang Cheng, Andrew Tsourkas, Robert Gorman, Victor A. Ferrari, Meagan McManus, Robert H. Mach, Joel S. Karp, Cory Tschabrunn, Giovanni Ferrari, Walter R. Witschey, Paco E. Bravo

**Author notes:** **Corresponding author contact information:** Paco E. Bravo, MD Divisions of Nuclear Medicine, Cardiothoracic Imaging and Cardiovascular Medicine Hospital of the University of Pennsylvania 3400 Civic Center Blvd, 11-154 South Pavilion Philadelphia, PA 19104. Drs. Witschey and Bravo are co-senior authors.

## Abstract

**Background:** Reactive oxygen species (ROS) contribute to myocardial ischemia-reperfusion injury (IRI), but in-vivo data on the spatial myocardial distribution and systemic effects of ROS after IRI remain limited. This multimodal CMR and PET/CT study aimed to non-invasively image ROS activity in a clinically-relevant swine model of IRI using [18F]ROStrace, a fluorine-18-labeled analogue of dihydroethidium (DHE), and to investigate regional changes in ROS activity in the infarcted myocardium during the subacute post-IRI phase.

**Methods:** IRI was induced by percutaneous occlusion of the left anterior descending artery for 90 minutes in swine (N=9). CMR and whole-body PET/CT imaging with [18F]ROStrace were performed before myocardial infarction (MI) and 3-5 days post-MI to assess ROS in non-infarct myocardium, lungs, bone marrow, spleen and skeletal muscle. Late gadolinium enhanced CMR was performed to structurally characterize infarct regions. Post-MI, in vivo [18F]ROStrace signal in infarcted myocardium was compared with remote, non-infarcted myocardium and validated via ex vivo DHE fluorescent imaging. Bulk RNA-sequencing (RNA-seq) and Gene Ontology pathway analysis were conducted on biopsies from infarct and remote myocardial tissue to identify differentially expressed genes and pathways connected to oxidative stress.

**Results:** During the subacute phase following MI, [18F]ROStrace fractional uptake rate (FUR; min-1) was significantly increased in skeletal muscle, compared to baseline (0.011±0.003 vs 0.016±0.005, p=0.04), with a trend toward increased FUR in bone marrow (0.046±0.009 vs 0.056±0.011, p=0.12) and the left ventricular free wall (0.067±0.007 vs 0.073±0.010, p=0.15). Within the myocardium, [18F]ROStrace FUR ((min-1)/(mL/min/g)) was significantly higher in infarcted compared to non-infarcted myocardium regions (0.110±0.034, vs 0.148±0.035, p=0.0005). DHE staining confirmed elevated ROS levels in the infarcted myocardium. RNA-seq identified 8,707 differentially expressed genes between infarct and remote myocardium, with downregulated pathways in the infarct associated with mitochondrial function, cellular respiration, and metabolic adaptation.

**Conclusion:** This study demonstrated MI ROS imaging using [18F]ROStrace using a whole-body PET/CT scanner and structural assessment with CMR. Systemic and myocardial increases in ROS activity were observed post-MI, accompanied by substantial molecular alterations in infarcted tissue. These findings show potential imaging strategies to evaluate therapeutic targets that can mitigate oxidative stress after MI.

## INTRODUCTION

Ischemic heart disease is the leading cause of death worldwide, accounting for nearly nine million deaths in 2017, and the prevalence of ischemic heart disease is predicted to increase from 1,655 per 100,000 individuals in 2017 to 1,845 per 100,000 by 2030 ^1^. Timely revascularization is critical for the reduction of myocardial infarction (MI) size, morbidity and mortality. While primary percutaneous coronary revascularization has improved outcomes for patients over the last 30 years, 30-day and 1-year mortality remains substantial at around 8% and 13%, respectively ^2,3^. Paradoxically, reperfusion can itself induce further injury in up to 40% of patients and has been associated with increased risk of arrhythmia and adverse left ventricular (LV) remodeling ^4^. Additionally, there may be molecular changes in extra-cardiac tissues after acute MI indicative of a systemic inflammatory response, including in the brain and hematopoietic tissues ^5^.

The pathophysiology of ischemic reperfusion injury (IRI) is still under investigation, but it is well established that reperfusion leads to increased reactive oxygen species (ROS) formation in cardiomyocytes ^6^. ROS are short-lived, highly reactive species that are derivatives of reduced O_2_, including superoxide, hydrogen peroxide, and hydroxyl radicals. ROS are created in healthy cells as a result of electron leakage during oxidative phosphorylation and play roles in cell signaling and in modulating the physiological response to stress in the cell ^7^. However, in reperfusion injury, the production of ROS far exceeds the antioxidant capability of the tissue, and excessive ROS production (i.e. oxidative stress) can cause single-strand DNA breaks, as well as lipid peroxidation and protein oxidation which damage lipid membranes and proteins. Calcium overload during reperfusion also opens the mitochondrial permeability transition pore, resulting in mitochondrial energy dysfunction and further ROS production ^8^.

Multiple technologies have been developed for detecting ROS in cell cultures and tissue sections. These include chemiluminescence assays, electron spin resonance, and fluorescence-based techniques with oxidizable fluorescent probes such as Peroxy Green 1, Peroxy Crimson 1, 2,7-dichlorodihydrogluorescein diacetate, dihydrorhodamine 123, hydrocyanines, and dihydroethidium (DHE) and its analogues ^9–11^. However, these techniques are unable to detect ROS in the myocardium *in vivo*. Positron emission tomography (PET) is a molecular imaging method that provides sensitive detection of radiotracer uptake which, when paired with the high spatial resolution provided by cardiac magnetic resonance (CMR) imaging, can enable localized detection of molecular probes *in vivo*.

[18F]ROStrace is a DHE-analog radiotracer sensitive to ROS, which has been shown to image elevated ROS in preclinical models of neuroinflammation ^12^. Moreover, newer multi-ring PET systems, such as our PennPET Explorer PET/CT scanner with a long axial field-of-view of 142 cm, can enable high-sensitivity dynamic imaging of the major body organs in large animals and humans ^13^, which allows for simultaneous acquisition of [18F]ROStrace data in organs that cannot be imaged simultaneously (e.g. brain and heart) with standard axial field-of-view (20 – 24 cm) PET/CT systems.

In this study, we measured ROS production in cardiac and extra-cardiac tissues in a swine model of subacute reperfusion injury *in vivo* using whole-body [18F]ROStrace PET/CT imaging and cardiac magnetic resonance (CMR). We hypothesized that [18F]ROStrace PET would detect sustained myocardial and systemic ROS activity during the subacute phase of ischemia-reperfusion injury and that these findings would correlate with ex vivo fluorescence and transcriptomic profiles.

## MATERIALS AND METHODS

### Experimental design and animal procedure

Healthy Yorkshire swine (N = 9) were included in this study, of which seven underwent baseline whole-body PET/CT imaging prior to myocardial infarction (MI) induction. Ischemia-reperfusion injury (IRI) was then produced by balloon occlusion of the mid–left anterior descending (LAD) coronary artery for 90 minutes, with angiographic confirmation of balloon placement immediately distal to the first diagonal branch. Acute myocardial injury was verified by ST-segment elevation on electrocardiography (ECG). Three to seven days after reperfusion, all nine animals underwent sequential cardiac magnetic resonance (CMR) and PET imaging on the same day.

Two days post-multimodal imaging, a terminal procedure was conducted and the heart was removed (**Figure 1**). Animals were under anesthesia with isoflurane for all imaging and terminal procedures. This study was approved by the Institutional Animal Care and Use Committee of the University of Pennsylvania.

**Figure 1:**
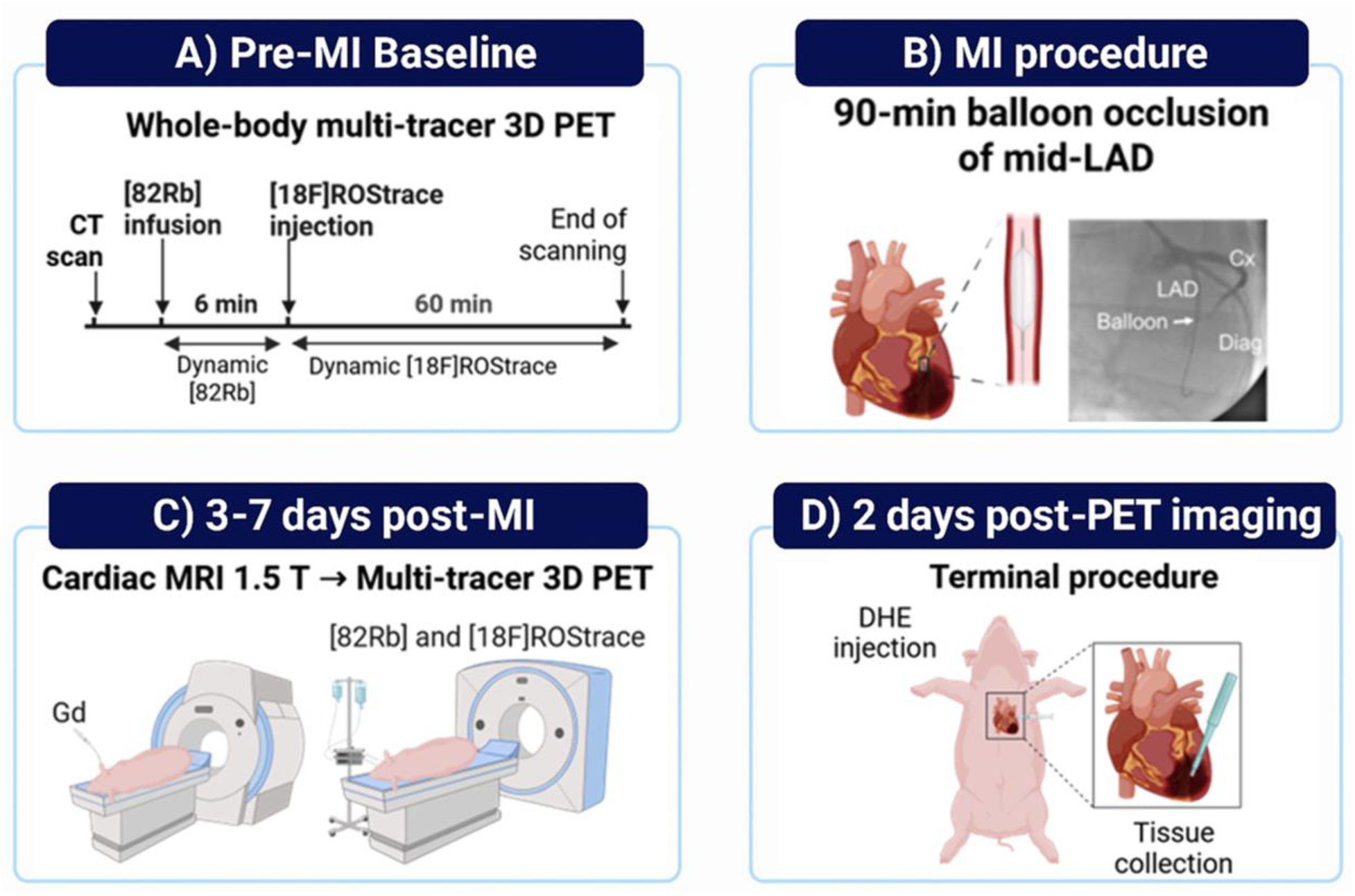
Schematic of the experimental setup. A) A whole-body baseline [82Rb] and [18F]ROStrace PET scan was performed in N=7 pigs. B) A 90-minute ischemia reperfusion injury was performed in N=9 pigs. C) All animals were scanned with CMR, including late gadolinium (Gd) imaging, and whole-body PET 3-7 days post-infarction. D) Tissue collection occured at 10 days post-MI and samples from N=2 pigs were analyzed. Created in Biorender.

### CMR protocol

CMR imaging was conducted on a 1.5 T whole-body system (Avanto, Siemens Healthcare, Erlangen, Germany). Cardiac gating was performed using a 3-lead ECG. Respiratory gating was achieved by inducing apnea at end-expiration by temporarily turning off the animal respirator. Short-axis cine images and cardiac and respiratory-gated late gadolinium enhanced (LGE) images were collected (technical details and image analysis in Supplementary Methods).

### Whole-body PET imaging

Animals were imaged on the long AFOV PennPET Explorer whole-body PET/CT system using a multi-tracer protocol that included 82-Rubidium [82Rb] for quantifying myocardial blood flow (MBF) and [18F]ROStrace for quantification of cardiac and systemic ROS levels in tissue.

### Radiotracer synthesis

[18F]ROStrace was synthesized in our cyclotron facility as previously described ^14^, which is near the PennPET Explorer PET/CT scanner (5-minute walk).

[82Rb] was produced from a standard 82-Strontium / 82Rb generator elution system developed by Bracco Diagnostics (CARDIOGEN-82). The lifespan of the generator ranged between 5-8 weeks for pre-clinical use only.

### Multi-tracer PET imaging protocol

Swine were positioned with the help of a CT topogram to include the whole body in the scanner FOV. Then, a CT scan (120 kV, 40 mA) mirroring the same FOV was obtained for attenuation correction of the PET data, ∼10 mCi (370 MBq) of [82Rb] was infused intravenously, and 3-dimensional list-mode scans were acquired for 6 minutes and reconstructed to static (2-min prescan delay) and 23-frame-dynamic images (14 frames x 5-sec, 5 frames x 10-sec, and 4 frames x 120-sec). Following [82Rb] scanning, ∼5 mCi (185 MBq) of [18F]ROStrace was administered intravenously and a 3-dimensional list-mode scan acquired for 60 minutes, and re-sampled to static (15 to 60 min) and 33-frame dynamic images (12 frames x 10-sec, 8 frames x 15-sec, 3 frames x 60-sec, 4 frames x 120-sec, 3 frames x 300-sec, 3 frames x 600 sec).

### Whole body PET image analysis

Whole body attenuation-corrected PET images were reconstructed and segmented using a commercial software package (MIMvista Corp, Cleveland, OH, USA). On baseline and post-MI images, 3D volumes of interest (VOIs) were drawn on delayed time frames in the LV free wall (a region expected to be spared from direct injury post-MI), vertebral bone marrow, whole brain, lung, quadriceps femoris skeletal muscle, and the spleen. The liver and kidney were not analyzed due to potential confounding effects of radiotracer excretion. The blood pool [18F]ROStrace signal was measured in the LV cavity. VOIs were subsequently translated onto all dynamic frames to generate tissue-specific time activity curves (TAC), and [18F]ROStrace fractional uptake rate (FUR) [min^-1^] was calculated as follows:

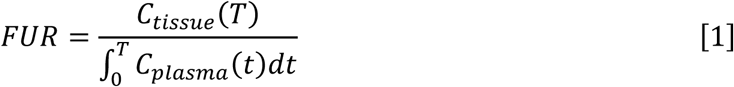

where 𝐶_𝑡𝑖𝑠𝑠𝑢𝑒_(𝑇) is the tracer concentration in the tissue at time=T, measured at equilibrium, and 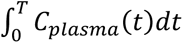 is the arterial input function.

### Cardiac PET image analysis

Attenuation-corrected static and dynamic [82Rb] and [18F]ROStrace PET datasets were volumetrically sampled, and LV polar maps were generated using dedicated cardiac analysis software (Carimas, Turku PET Centre, Finland). Infarcted and remote myocardial regions were defined directly from static [82Rb] PET images summed between 2 and 6 minutes post-injection. The infarct region was identified using Otsu’s method for histogram-based thresholding, which automatically determines the optimal cutoff separating low- and high-intensity voxels within the LV polar map. Areas below the Otsu-derived threshold corresponded to infarcted myocardium, which consistently involved the apex and mid-to-distal anteroseptal segments (mid-LAD vascular distribution). The LV free wall, which was spared from infarction in all cases, was designated as the remote (non-infarct) region. Myocardial blood flow (MBF, mL/min/g) was quantified using a standard one-compartment kinetic model, and both uncorrected and MBF-corrected fractional uptake rate (FUR) values for [18F]ROStrace were extracted for infarct and remote myocardial regions.

### Imaging statistical analysis

Shapiro-Wilk tests were performed to assess normality of the calculated FUR for each tissue. Changes in [18F]ROStrace fractional uptake rate between baseline and post-MI in the brain, bone marrow, lung, LV free wall, spleen, and skeletal muscle were evaluated using paired t-tests for each tissue, and Cohen’s d effect sizes (ES) were calculated. Paired t-tests were used to evaluate the difference in MBF-corrected [18F]ROStrace FUR between regions of infarct and non-infarct myocardium post-subacute MI.

### Dihydroethidium (DHE)

Ex-vivo experiments with in-vivo DHE injection were conducted in two animals. Preparation of the DHE solution is detailed in the Supplementary Methods. DHE solution was prepared for left and right selective heart coronary injections, and DHE injection was performed in the left coronary followed by the right coronary system while the animal was still alive. The heart was arrested one-hour post-DHE injection with an infusion of cold cardioplegia solution and harvested. Heart tissue biopsies from the LV infarct region and free wall were collected and fixed in formalin. Light-protected paraffin blocks were sectioned into 5µm slices. Sections were also stained with Masson Trichrome for fibrosis.

Images were collected with a Nikon AX confocal microscope system (561 excitation, red emission. Obj 20X) and quantified using ImageJ software. Quantification of fluorescence intensity of DHE was calculated by measuring the corrected total cell fluorescence parameter: = Integrated Density – (area x mean fluorescence of background readings).

DHE statistical analysis: Results are the average ± SD of a total of 6 LV free wall areas and 10 LV infarct areas (pig N=2). A Shapiro-Wilk normality test was performed and a t-test was used to determined significant data with p<0.05 (*). Statistical tests were performed using GraphPad.

### Bulk RNA-sequencing analysis

mRNA was isolated from post-MI infarct and non-infarct free wall biopsy samples for RNA sequencing (Supplementary Methods). RNA sequencing was performed by the Columbia genomics core. Bulk RNA-seq raw fastq files were first subjected to quality control using FastQC^15^ (v0.12.1) to identify potential issues such as contamination and adapter sequences. After trimming low-quality sequences, FastQC was rerun on the cleaned reads to ensure that all major quality issues were resolved. The sequencing reads were then aligned to the Sus scrofa reference genome (Sscrofa11.1) using HISAT2 (v2.2.1). Gene expression levels were quantified with featureCounts (v2.0.6), and differential expression analysis was conducted using DESeq2. **Transcriptomic statistical analysis**

The differentially expressed genes (DEGs) between different conditions were determined using the Wald test with a cut-off for absolute log2 fold change greater than 0.5 and adjusted p-values less than 0.05. Gene ontology (GO) analysis was conducted by using the ShinyGO application developed based on several R/Bioconductor functions and Enrichr database. Finally, volcano plots and GO pathways were generated with the ggplot2 package in R.

## RESULTS

### *In vivo* imaging of ROS

We first sought to determine the systemic biodistribution of [18F]ROStrace at baseline and after IRI. [18F]ROStrace signal was observed in multiple organs, including the heart, brain, bone marrow, lung, skeletal muscle and spleen (**Figure 2A**). Compared to baseline, [18F]ROStrace FUR increased significantly in skeletal muscle (0.011±0.003 vs 0.016±0.005; *p*=0.04) post-MI, and there was trend of increased ROS in bone marrow (0.046±0.009 vs 0.056±0.011; *p*=0.12) and LV free wall (0.067±0.007 vs 0.073±0.010; p=0.15). By contrast, brain (0.051±0.007 vs 0.050±0.004; p=0.68), lungs (0.022±0.004 vs 0.019±0.004; p=0.14), and spleen (0.021±0.003 vs 0.020±0.002; p=0.31) FUR did not change (**Figure 2B, 2C**).

**Figure 2:**
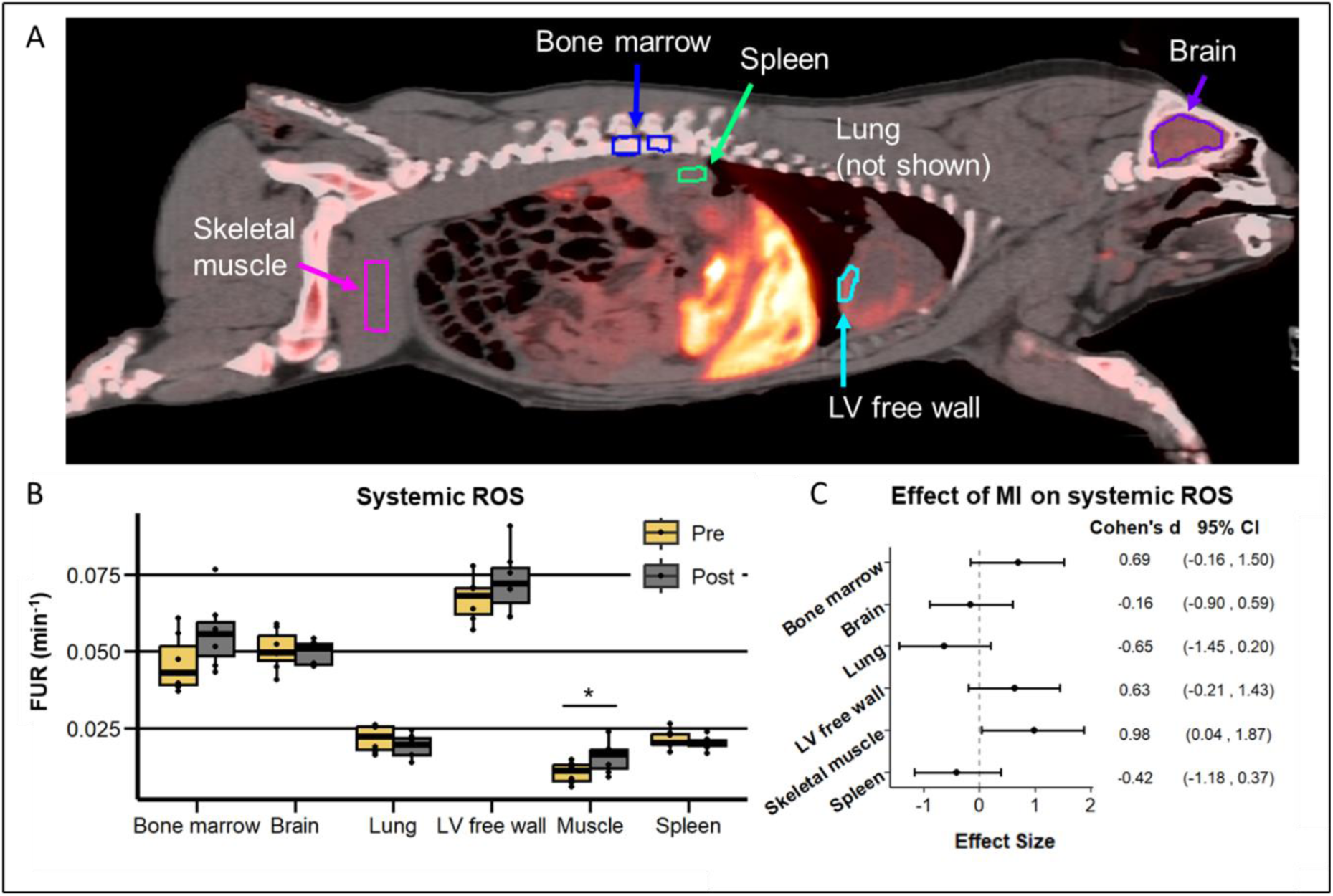
Whole-body imaging of [18F]ROStrace. **A)** [18F]ROStrace image (summed frames 15-60 minutes) in the sagittal plane showing baseline ROS in the heart and extra-cardiac tissue. **B)** [18F]ROStrace uptake in extra-cardiac tissue at baseline and post-MI. **C)** Cohen’s D effect size, indicating the strength of the effect of MI on [18F]ROStrace uptake. ROS was increased in skeletal muscle and trended toward a moderate increase in bone marrow and LV free wall post-MI; Paired T-test p<0.05 (*).

### CMR characterization of infarction

Left ventricular functional measurements derived from CMR post-infarct are reported in **Supplemental Table 1**. Infarct size measured on LGE was 16.2 ± 2.3 g, MVO size was 2.2 ± 2.0 g, and ejection fraction was 31.5 ± 7.8 %.

### *In vivo* measurement of ROS in post-IRI myocardium

We next assessed whether there was a change in ROS levels post-MI between infarcted myocardium in the anteroseptal wall and apex versus non-infarct myocardium in the LV free wall that is detectable with [18F]ROStrace PET imaging. First, the time activity curves of [82Rb] and [18F]ROStrace demonstrated that both tracers exhibit relative sustained level of activity after blood pool clearance in the infarct and non-infarct tissue (**Figure 3**). MBF-corrected [18F]ROStrace [(min^-1^)/(mL/min/g)] was significantly higher in the infarcted myocardium region than in the remote non-infarcted LV free wall (0.110±0.034, vs. 0.148±0.035; P=0.0005) (**Figure 4A, B**). As a preliminary validation, [18F]ROStrace imaging was compared to DHE imaging (**Figure 4C**). Analysis of pig heart biopsies showed increased DHE accumulation in the infarct area compared to the remote non-infarct LV free wall tissue (**Figure 4D**), supporting the [18F]ROStrace measurement of ROS *in vivo*.

**Figure 3:**
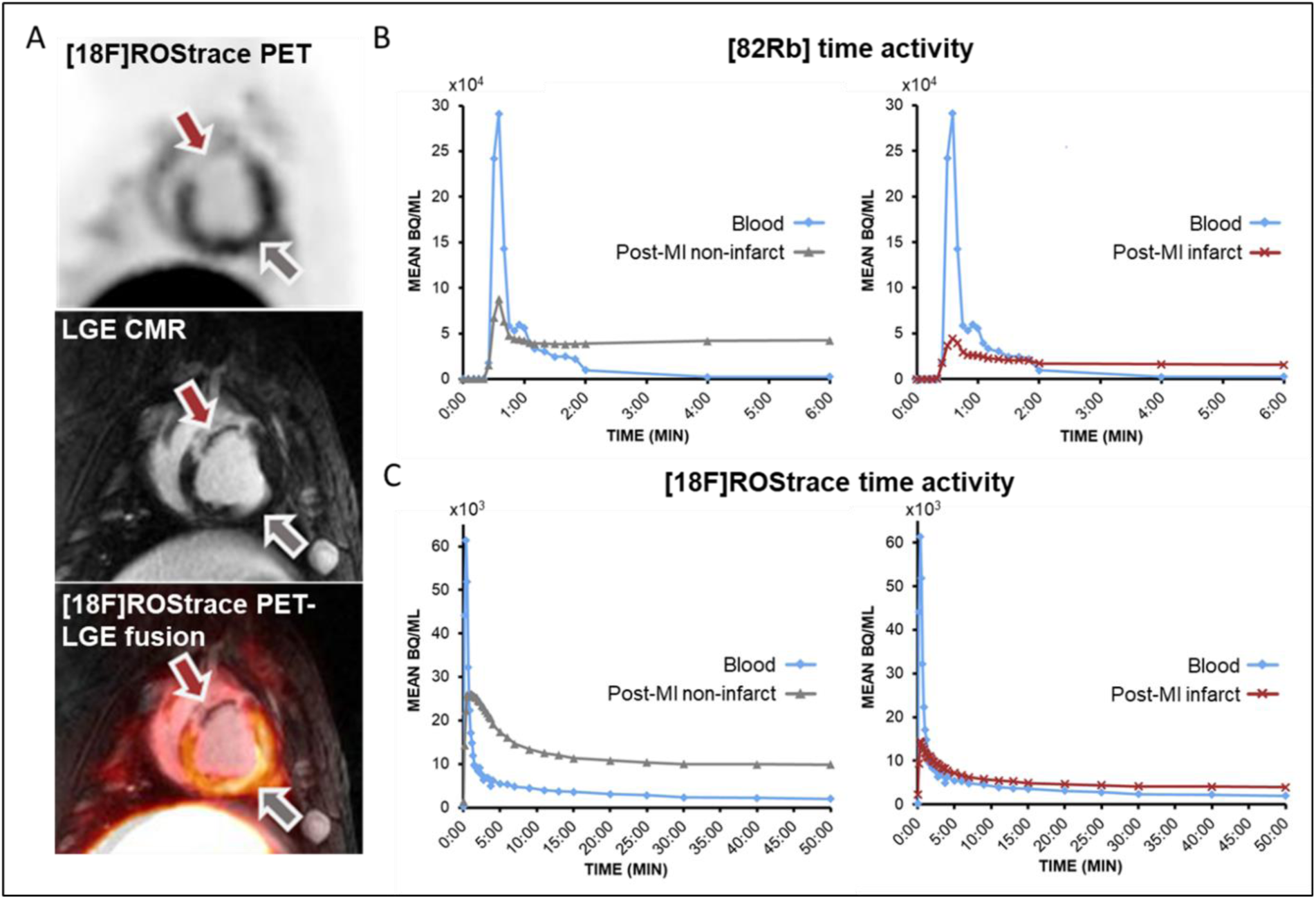
*In vivo* imaging of myocardial ROS. **A)** Short axis [18F]ROStrace (top, summed frames 15-60 minutes), late gadolinium enhanced MR (middle) in the same animal, and PET-MR overlay after MI (bottom) showing the region of infarct (red arrow) and non-infarct free wall myocardium (gray arrow). Time-activity curves of **B)** [82Rb] and **C)** [18F]ROStrace uptake in the myocardium in infarct (red lines) and non-infarct post-MI (gray lines), as well as in the arterial blood pool (blue lines) for determination of the arterial input function (AIF).

**Figure 4:**
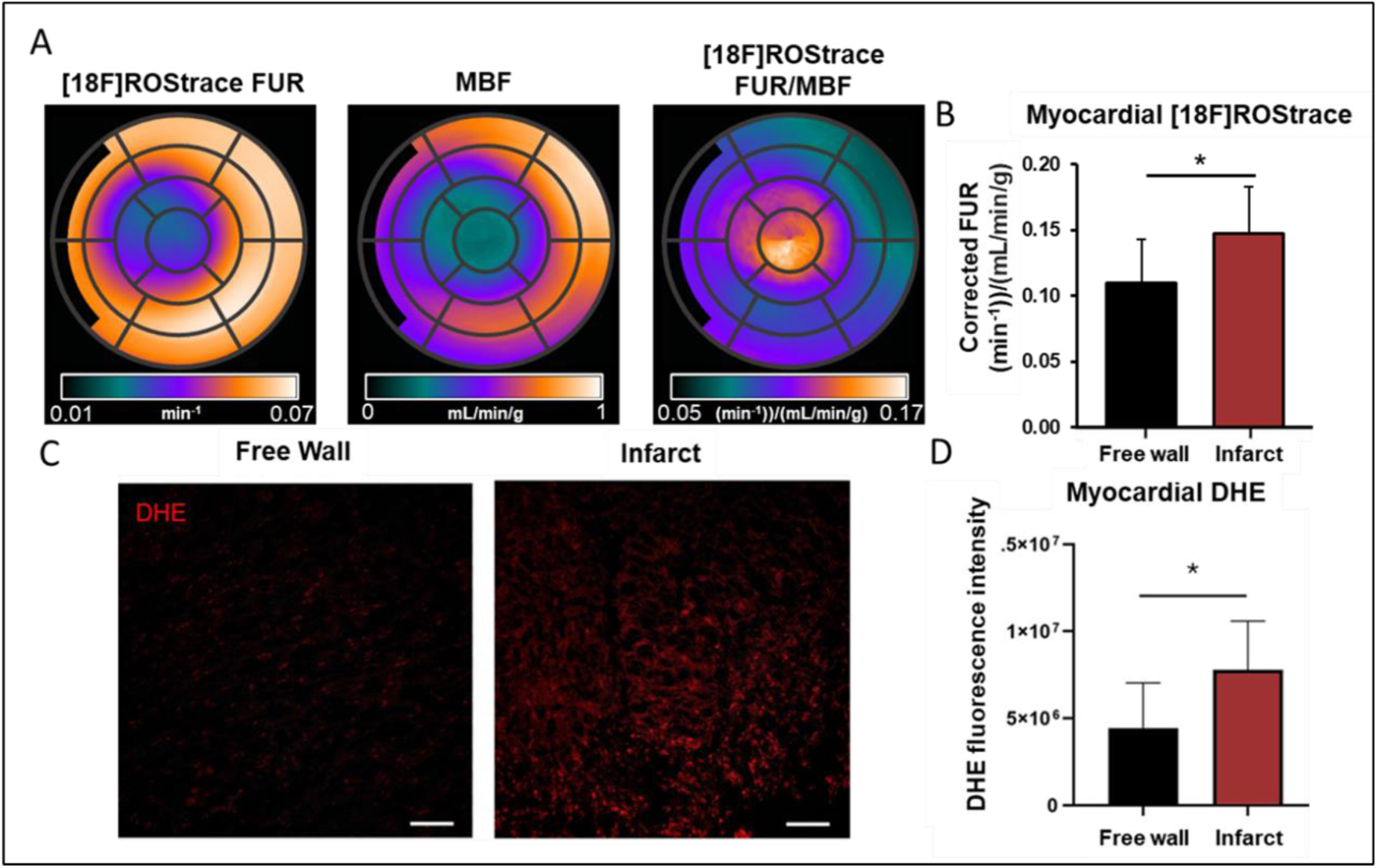
ROS is increased in infarcted myocardium. **A)** AHA 17-segment polar maps showing the absolute [18F]ROStrace, FUR, MBF, and the [18F]ROStrace FUR corrected by MBF. **B)** Corrected [18F]ROStrace FUR is increased in infarct myocardium compared to remote free wall myocardium; Paired T-test p<0.05(*). **C)** Infarct and free wall tissue biopsies of a heart injected with DHE before arrest. Formalin-fixed slides imaged with confocal microscopy (scale bar=100µm). **D)** DHE (*red*) fluorescence intensity is measured as corrected total cell fluorescence (CTCF)/area. Statistical analysis: results are the average ± SD of a total of 6 (free wall) areas and 10 (infarct) areas; T-test p<0.05 (*).

### Differentially expressed gene (DEG) analysis in the heart post-MI

To further investigate the alteration in ROS after IRI, we evaluated global gene expression differences between the remote free wall and infarct area using bulk RNA-seq analysis and Gene Ontology (GO) pathway analysis. After removing one identified outlier, principal component analysis (PCA) of the bulk RNA-seq data showed that sample groups were highly homogeneous and highly separated (**Supplemental Figure 1**).

Our DEG analysis identified a total of 8,707 DEGs when comparing free wall versus infarct heart areas. Among these DEGs, 4506 were upregulated and 4201 were downregulated in the free wall relative to the infarct (padj <0.05 & |log2FC| >0.5). **Figure 5A** shows a volcano plot displaying the distribution of DEGs and highlighting the top 10 upregulated and downregulated genes. A heatmap (**Figure 5B**) examines the distinct gene expression profiles between the free wall and infarct among all the samples.

**Figure 5:**
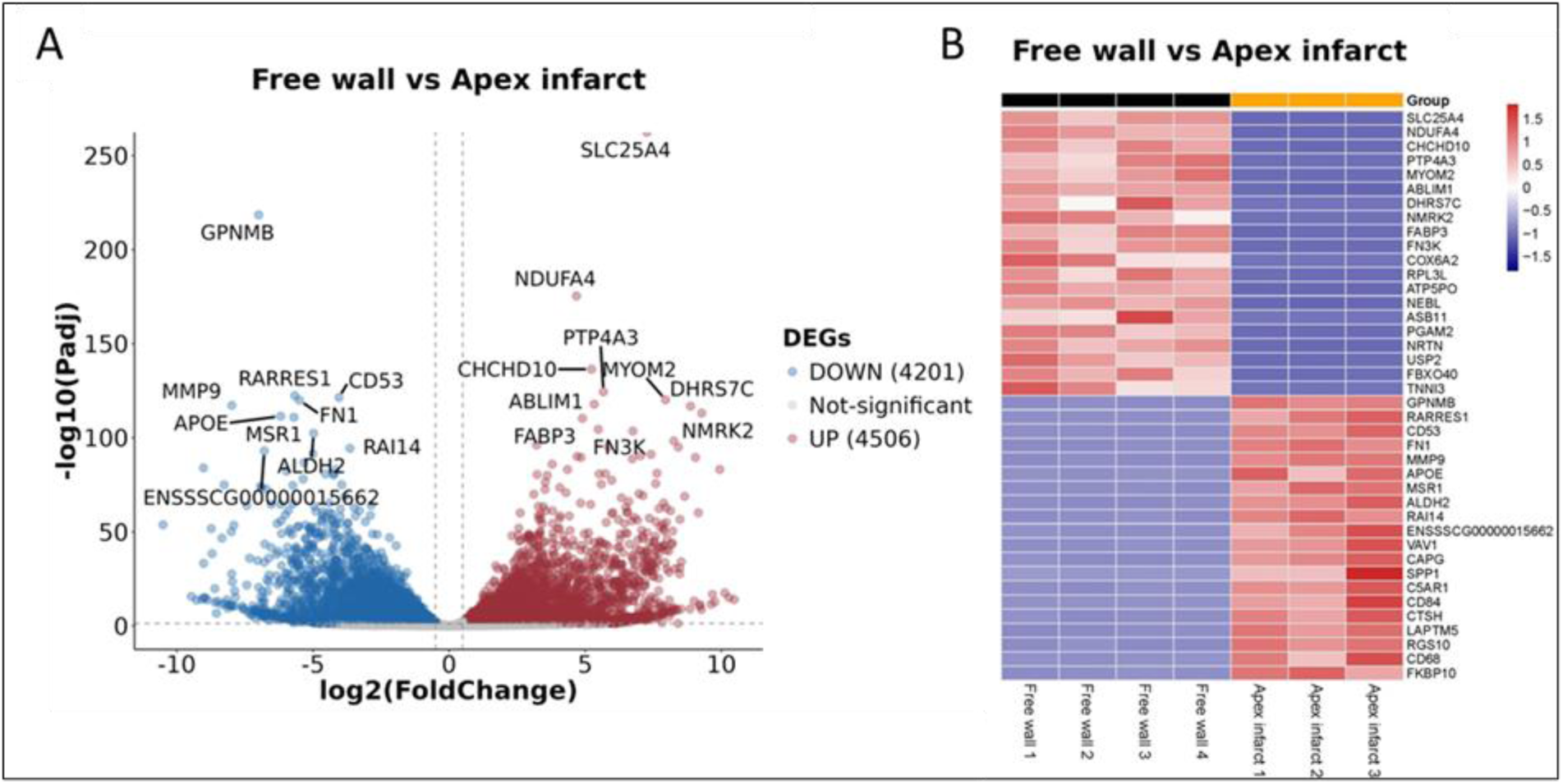
Differentially expressed genes – DEGs: comparison between free wall vs. apex infarct tissue biopsies. **A)** Volcano plot showing the DEGs. Genes with log 2-fold change of p >0.5 and padj <0.05 were considered differentially expressed. Blue dots are the down-regulated DEGs in the free wall; red dots are the up-regulated DEGs in free wall; grey dots are not significant according to the padj value. **B)** Heatmap of genes with log 2-fold changes >0.5 in the comparisons free wall (*black*) vs infarct (*yellow*).

To better characterize the statistically significant DEG function, we compared the listed genes with Enrichr database gene sets available related to ROS. In the infarct, several DEGs emerged as downregulated in the oxidative phosphorylation, mitochondrial respiration, and cardiac protection against ROS gene sets. Upregulated DEGs were highlighted in the oxidative stress and extracellular matrix remodeling gene sets (**Supplemental Table 2**).

### Functional and pathway analysis

We further interrogated bulk RNA-seq data to investigate the mechanisms associated with the occurrence and development of MI. GO enrichment pathway analysis of the DEGs showed that in the infarct, at the biological process level, these genes were predominantly downregulated in pathways related to mitochondrion organization, cellular respiration, aerobic respiration, mitochondrial respiratory chain complex assembly, oxidative phosphorylation, and heart contraction. Furthermore, KEGG pathway analysis suggested that the identified DEGs were significantly involved in pathways related to oxidative phosphorylation, metabolic pathways and cardiac muscle contraction (**Figure 6A**). **Figure 6A** also shows the GO cellular component pathway network of DEGs in the infarct vs free wall. One main downregulated hub in the network was mitochondrion and all the pathways related to it, such as mitochondrial respirasome, oxidoreductase complex, respiratory chain complex, and mitochondrial protein-containing complex. The downregulation of these biological processes is involved in mitochondrial activity and ROS production. Among the upregulated biological processes (**Figure 6B**), pathways related to metabolic adaptation, cell transportation, and stress response were prominently observed in the infarct.

**Figure 6:**
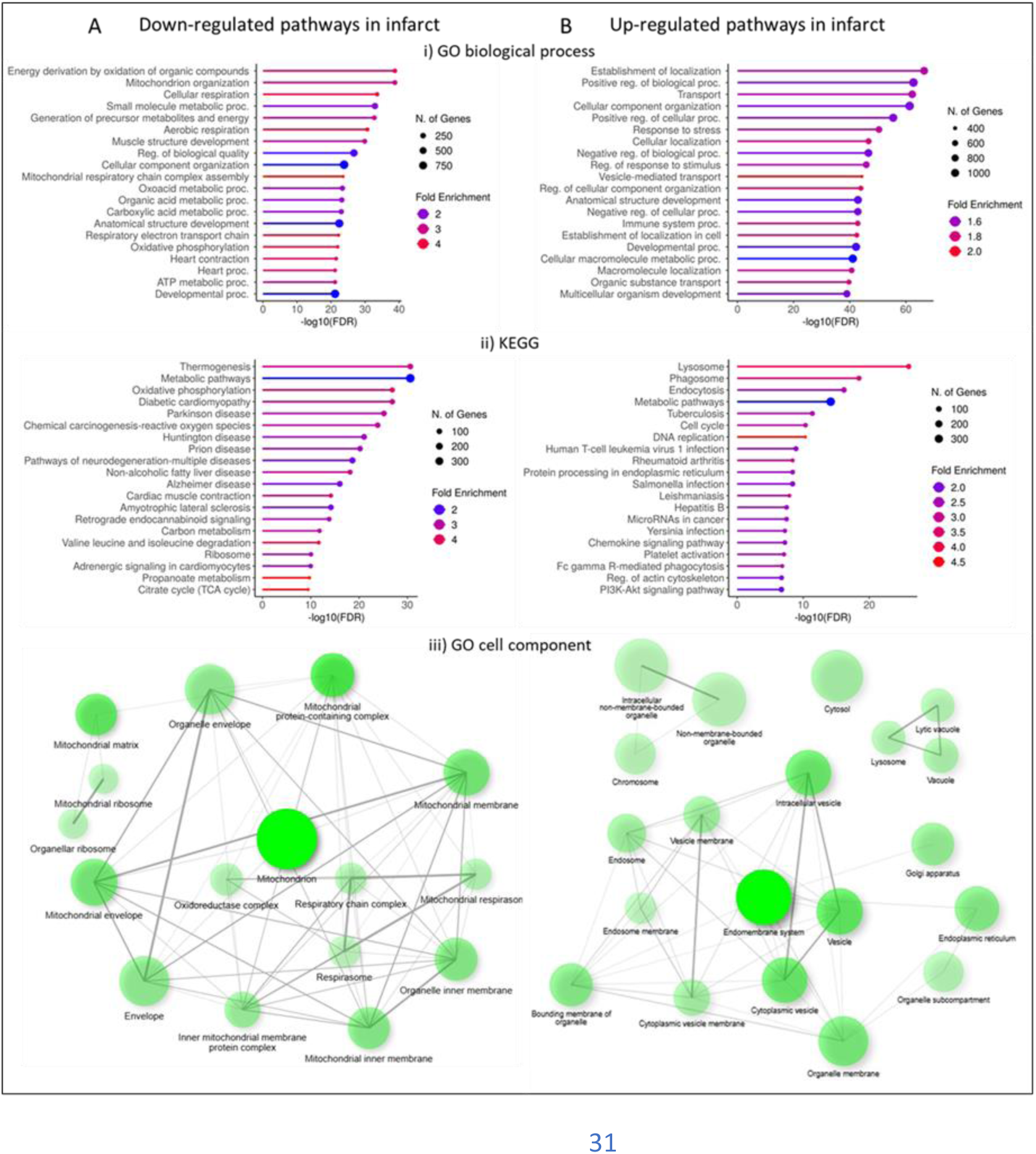
Enriched pathways among the genes expressed differentially between free wall and infarct areas. **A)** Down-regulated pathways in apex infarct tissues. **B)** Up-regulated pathways in apex infarct tissues. **i)** GO pathway biological process and **ii)** KEGG (Kyoto Encyclopedia of Genes and Genomes) database. FDR (False Discovery Rate) = Padj value (<0.05). The 20 top-ranked Genomes pathways are listed. **iii)** GO pathway cellular component database. Network graphical representation shows that two pathways (nodes) are connected if they share 20% (default) or more genes. Darker nodes are more significantly enriched gene sets. Bigger nodes represent larger gene sets. Thicker edges represent more overlapped genes.

## DISCUSSION

In this study, we demonstrated the feasibility of imaging myocardial and systemic ROS in vivo using [18F]ROStrace PET combined with CMR in a large-animal model of ischemia-reperfusion injury. ROS activity was significantly increased in infarcted compared with remote myocardium during the subacute phase after MI, and these findings were validated by ex vivo DHE fluorescence. Transcriptomic analysis revealed broad alterations in mitochondrial and oxidative stress pathways, further supporting a persistent oxidative phenotype within the infarcted myocardium. Together, these data provide a foundation for applying molecular PET imaging to study oxidative stress and evaluate ROS-targeted therapies after reperfusion.

### *In vivo* imaging of ROS before and after MI

This work represents the first use of PET to assess ROS in vivo in a large-animal model of IRI with multimodal validation. ROS play a central role in reperfusion injury by disrupting cellular membranes, proteins, and mitochondrial function^6–8,16^. Quantification of ROS in vivo has been challenging because of their short half-lives and the limitations of existing biochemical or fluorescence-based assays^9–11^. [18F]ROStrace, a dihydroethidium (DHE) analog sensitive to superoxide, enables quantitative imaging of oxidative stress in living tissue ^12,14,17,18^. In our study, whole-body PET imaging revealed both myocardial and extra-cardiac ROS activity, extending prior small-animal data on DHE-analog tracers.

ROS increases were most pronounced in infarcted myocardium, with additional trends in skeletal muscle and bone marrow. The skeletal muscle findings are consistent with previous reports of oxidative stress and altered oxygen metabolism after MI^19,20^, whereas increased bone marrow signal may reflect inflammatory activation and impaired release of endothelial progenitor cells ^21^. In contrast to previous studies reporting post-MI brain uptake of other inflammatory biomarkers such as the mitochondrial translocator protein^5,22^, no significant change was observed with [18F]ROStrace, suggesting that systemic oxidative alterations during the subacute phase are tissue-specific.

### ROS activity in the subacute infarcted myocardium

Prior studies in cell culture and small animals have demonstrated a burst of ROS generation during ischemia and early reperfusion^6,23^, but the persistence of oxidative stress over subsequent days has remained uncertain. Our results show sustained elevation of ROS within the infarct zone three to seven days after reperfusion, suggesting ongoing oxidative injury and mitochondrial dysfunction. This observation aligns with human data linking persistent redox imbalance after MI with adverse remodeling and late cardiac events ^24^. Because infarcted regions have reduced perfusion and viable tissue, we corrected [18F]ROStrace uptake for myocardial blood flow, confirming that the signal represents true biochemical activity rather than delivery differences.

In a subset of animals, coronary injection of DHE confirmed increased superoxide activity in infarcted myocardium, supporting the in vivo PET findings. While DHE staining in large-animal models is technically challenging^25^, the concordant increase in both [18F]ROStrace uptake and DHE fluorescence suggests that the PET signal reflects sustained oxidative stress rather than a methodological artifact. Persistent ROS activity beyond the acute phase may perpetuate injury and contribute to progressive ventricular remodeling.

### Molecular pathways associated with ROS after IRI

Bulk RNA-sequencing analysis identified 8,707 differentially expressed genes between infarct and remote myocardium, indicating broad transcriptional remodeling after reperfusion. Downregulated pathways were enriched for mitochondrial respiration, oxidative phosphorylation, and contractile function, consistent with impaired energy metabolism^26^. Upregulated pathways involved cellular stress responses, extracellular matrix remodeling, and metabolic adaptation. Notably, superoxide dismutase (SOD1, SOD2) and mitochondrial complex I–IV components (NDUFA4, NDUFS4, NDUFV1) were suppressed in infarcted tissue, suggesting reduced antioxidant defense and mitochondrial dysfunction as contributors to sustained ROS generation^27–30^. These molecular alterations align with prior evidence that ROS-mediated mitochondrial injury contributes to both cardiomyocyte loss and fibrotic remodeling. Extracellular matrix remodeling post-MI, including upregulation of MMP9 and TGFβ, may further amplify oxidative and inflammatory stress^31,32^. These findings highlight mitochondrial and extracellular matrix pathways as key determinants of oxidative injury and repair after IRI.

### Translational significance

Persistent oxidative stress after MI has been linked to adverse remodeling, arrhythmogenesis, and progression to heart failure^4,32^. The ability to quantify ROS noninvasively could identify patients with elevated oxidative burden and facilitate evaluation of emerging redox-targeted therapies. Although antioxidant interventions such as superoxide dismutase administration have yielded mixed results in reducing infarct size, they have been associated with reductions in reperfusion arrhythmias^33^. More recent therapeutic approaches, including ROS-modulating nanoparticles and hydrogel-based targeted delivery systems, require sensitive imaging endpoints to assess efficacy^34^. [18F]ROStrace PET offers a translationally feasible method for noninvasive quantification of ROS across multiple organs, potentially guiding the development and evaluation of cardioprotective therapies.

### Study Limitations

There were several limitations in this study. First, the sample size was relatively small but consistent with other large-animal models, limiting the statistical power to detect differences in ROS production at baseline versus post-infarction across all organs. In addition, while the induced coronary occlusion produced a reproducible injury in terms of location and severity, it does not capture the heterogeneity observed in human myocardial infarction. All imaging was performed under anesthesia, which may influence oxidative metabolism and ROS generation. However, the observed increase in ROS activity within the infarct and selected extra-cardiac tissues warrants further investigation in larger cohorts. Finally, in vivo [¹⁸F]ROStrace PET imaging and ex vivo DHE fluorescence analyses were performed approximately two days apart. Despite this temporal separation, both modalities demonstrated concordant increases in ROS activity within the infarct region, suggesting that oxidative stress remains elevated during the subacute phase and that these findings reflect a sustained biological response rather than a transient or methodological artifact.

## CONCLUSION

This study establishes [18F]ROStrace PET as a feasible and biologically meaningful approach for imaging oxidative stress in the subacute phase after myocardial infarction. The integration of PET, CMR, and transcriptomics provides complementary insights into the molecular and structural remodeling that accompany reperfusion injury. Persistent ROS activation within infarcted myocardium and systemic tissues supports the concept of oxidative stress as a sustained and quantifiable therapeutic target. Future studies in larger cohorts and eventual human translation will be essential to validate [18F]ROStrace PET as a biomarker for oxidative injury and treatment response after reperfusion.

## Acknowledgements

We gratefully acknowledge the Penn cyclotron staff for radio-tracer production and PennPET Explorer staff for data acquisition and processing.

## Supplemental Material

Supplemental Methods

Tables S1–S2

Figure S1

## Non-Standard Abbreviations and Acronyms

MI: Myocardial infarction
ROS: Reactive oxygen species
IRI: Ischemic reperfusion injury
DHE: Dihydroethidium
PET: Positron emission tomography
DHE-analog PET radiotracer: ROStrace
CMR: Cardiac magnetic resonance
LAD: Left anterior descending coronary
82Rb: 82-Rubidium
MBF: Myocardial blood flow
AFOV: Axial field of view

