## Supplementary Methods for "In vivo imaging of reactive oxygen species after myocardial ischemia-reperfusion injury: a large animal multimodal imaging and transcriptomic study"

### CMR protocol

Cardiovascular magnetic resonance (CMR) imaging was conducted on a 1.5 T whole-body system using a transmit body coil and 6-channel body matrix receive coil integrated with a 9 to 12-channel spine matrix receive coil (Avanto; Siemens Healthcare). Cardiac gating was performed using a 3-lead ECG. Respiratory gating was achieved by inducing apnea at end-expiration by temporarily turning off the animal respirator. Retrospective, short-axis cine MRI was performed with parameters: [TR = 38.8-43.2 ms, TE = 0.94 ms, flip angle = 56°, FOV = 285 x 380 mm, spatial resolution = 2.97 x 2.97 $mm^2$, slice thickness = 3 mm, number of slices = 10-14, acceleration factor = 3, NEX = 1, cardiac phases = 15-20]. Cardiac and respiratory-gated late gadolinium enhanced (LGE) images were acquired using a 2D phase-sensitive inversion recovery (PSIR) segmented gradient echo sequence. LGE scan parameters were: TR= 535-750 ms, TE = 3.23 ms, flip angle = 25°, FOV = 201 x 340 mm, spatial resolution = 1.33 x 1.33 $mm^2$, slice thickness = 8 mm, number of slices = 12, acceleration factor = 2, NEX = 1.

### PET analysis and MBF quantification

Because uptake of ROStrace in myocardial tissue is affected by the amount of viable tissue, we corrected the [18F]ROStrace FUR for MBF in the infarct and non-infarct myocardial regions. Attenuation-corrected static and dynamic [82Rb] cardiac PET images were volumetrically sampled, and standard 17-American Heart Association (AHA) left ventricular (LV) polar maps were generated using commercially available software (Invia Medical Imaging Solutions, Ann Arbor, MI, USA). Infarct myocardium was segmented using Otsu's method for thresholding [73]. These infarct areas were identified in blackout polar maps and invariably corresponded to segments involving the apex and mid-distal anterior septum (mid-LAD vascular distribution). The free LV wall, on the other hand, was spared from infarction in all cases and was designated as the non-infarct / remote myocardium region. Subsequently, MBF (mL/min/g) was quantified at the infarct and non-infarct myocardial regions using a standard one-compartment model. In N=4 cases, model fitting failed and an MBF polar map could not be generated. For these cases, a MBF map created by averaging the MBF maps of the remaining cases was used. [18f]ROStrace FUR was corrected by MBF as follows:

$$Corrected\,[18F]ROStrace\,FUR = \frac{FUR_{ROStrace}}{MBF}. \qquad [1]$$

**DHE preparation**

DHE solid (Sigma Aldrich) was dissolved in DMSO (10 µL DMSO/mg DHE) under low-light conditions. The DHE/DMSO solution was then diluted to 1:100 using a 50/50 mixture of ethanol and Tween 20, and the resulting DHE/DMSO/EtOH/Tween solution was further diluted to 1:1000 with sterile saline for a final DHE solution concentration of 1 mg/mL. 2 mL and 1 mL of DHE solution was prepared for left and right selective heart coronary injections, respectively.

**RNA extraction and RNA integrity**

Isolation of total mRNA for RNA-seq was performed using the RNeasy mini kit (Qiagen, Valencia, CA). 1% β-mercaptoethanol was added to the lysis buffer and X mg of previously flash-frozen tissue was homogenized with a TissueRuptor (Qiagen). Isolated RNA purity was preliminarily assessed on a NanoDrop spectrophotometer (NanoDrop technologies). RNA concentration was measured on a Qubit 2.0 fluorometer (Invitrogen). RNA integrity assessment was performed on an Agilent Bioanalyzer 2100 using a Nano 6000 assay kit (Agilent Technologies) Columbia Genome Center core. Most of the samples presented an RNA integrity number (RIN) > 6.

**RNA sequencing**

A total of > 600 ng RNA per sample was used as input material for the RNA sample preparations. Sequencing libraries were generated using Illumina TruSeq Stranded mRNA kit and modified the protocol by replacing the Illumina TruSeq PCR reaction with KAPA HiFi HotStart Ready Mix for the final PCR step to make it compatible with the 2x75bp paired-end sequencing on the Aviti Element. RNA sequencing was performed by the Columbia genomics core.
