## Supplemental Materials for "In vivo imaging of reactive oxygen species after myocardial ischemia-reperfusion injury: a large animal multimodal imaging and transcriptomic study"

**Supplementary Material**

**Supplementary Table 1.** Summary of cardiac magnetic resonance measurements

| CMR functional measures | |
|---|---|
| Infarct Mass Size (g) | 16.2 ± 2.3 |
| MVO Mass Size (g) | 2.2 ± 2.0 |
| ED mass (g) | 74.3 ± 9.1 |
| EDV (ml) | 108.6 ± 11.9 |
| ESV (ml) | 74.7 ± 13.7 |
| EF (%) | 31.5 ± 7.8 |
| CO (l/min) | 3.2 ± 1.2 |

**Supplementary Table 2.** Enrichr gene sets containing differentially expressed genes found in post-MI myocardium in the subacute phase.

| | UP | DOWN | Fold Change | padj |
|---|---|---|---|---|
| **Oxidative phosphorylation** | | NDUFA4 | -4.6800 | 3.99E-176 |
| | | CoX6A2 | -6.7520 | 2.59E-104 |
| | | ATP5PO | -3.2134 | 5.757E-97 |
| | | COX7A1 | -7.1247 | 4.919E-70 |
| | | ATP5MC1 | -3.6059 | 8.37E-81 |
| | | PPARGC1A | -5.9738 | 3.478E-78 |
| | | NDUFS4 | -3.2813 | 2.932E-64 |
| | | NDUFV1 | -3.4683 | 3.864E-64 |
| | | COX8H | -9.1576 | 6.575E-61 |
| | | UQCRC1 | -3.2797 | 2.74E-59 |
| **Oxidative stress related genes** | MAPK14 | | 0.5600 | 0.033 |
| | CYBA | | 3.4466 | 2.69E-24 |
| | SOD3 | | 0.6950 | 0.015 |
| | GSR | | 1.9125 | 3.288E-12 |
| | GCLC | | 2.3445 | 2.979E-10 |
| | HMOX1 | | 5.5272 | 9.366E-54 |
| | APOE | | 6.1859 | 2.98E-112 |
| | | XDH | -7.1988 | 2.574E-24 |
| | | SOD2 | -1.1918 | 0.036 |
| | | SOD1 | -0.8545 | 0.006 |

| | UP | DOWN | Fold Change | padj |
|---|---|---|---|---|
| **Mitochondrial Respiration** | | ATP5PO | -3.2134 | 5.76E-97 |
| | | ATP5MC1 | -3.6059 | 8.37E-81 |
| | | CHCHD10 | -5.2349 | 3.23E-137 |
| | | ATP5MG | -2.8561 | 1.01E-38 |
| | | ATP5MK | -3.3735 | 2.77E-35 |
| | | ATP5IF1 | -2.8937 | 1.01E-32 |
| | | ATP5MC2 | -2.0100 | 1.92E-31 |
| | | ATP5ME | -4.2048 | 1.11E-29 |
| | | ATP5F1C | -2.8396 | 1.95E-29 |
| | | ATP5MC3 | -2.5924 | 1.25E-28 |
| | | ATP5PB | -2.6561 | 1.74E-20 |
| | | ATP5PF | -3.0812 | 3.71E-46 |
| | | ATP5F1D | -1.8783 | 4.5E-13 |
| | ATP6VOB | | 2.2204 | 4.50E-30 |
| | ATP6VOD2 | | 8.3477 | 2.1E-47 |
| **Extracellular matrix remodeling** | FN1 | | 5.5023 | 9.2E-121 |
| | MMP9 | | 7.9825 | 4.92E-118 |
| | SPP1 | | 9.0234 | 1.04E-84 |
| | ADAM12 | | 6.7053 | 2.25E-73 |
| | LGMN | | 3.8314 | 2.14E-69 |
| | MAFB | | 4.4052 | 3.52E-66 |
| | SDC1 | | 5.7339 | 8.88E-66 |
| | VMP1 | | 2.8633 | 7.59E-65 |
| | COTL1 | | 4.9413 | 2.31E-64 |

**Supplementary Figure 1.** Principal component analysis – PCA. The plot represents the distribution of the samples within the free wall *(blue)* and apex infarct *(red)* groups.

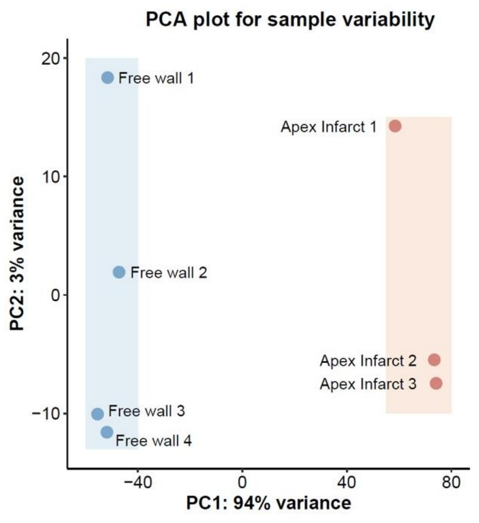
